# Dynamics of Livestock-Associated Methicillin Resistant *Staphylococcus aureus* in pig farms networks: insight from mathematical modeling and French data

**DOI:** 10.1101/727032

**Authors:** Jonathan Bastard, Mathieu Andraud, Claire Chauvin, Philippe Glaser, Lulla Opatowski, Laura Temime

## Abstract

Livestock-associated methicillin resistant *Staphylococcus aureus* (LA-MRSA) colonizes livestock animals worldwide, especially pigs and calves. Although frequently carried asymptomatically, LA-MRSA can cause severe infections in humans. It is therefore important to better understand LA-MRSA spreading dynamics within pig farms and over pig farms networks, and to compare different strategies of control and surveillance. For this purpose, we propose a stochastic meta-population model of LA-MRSA spread along the French pig-farm network (n=10,542 farms), combining within- and between-farms dynamics, based on detailed data on breeding practices and pig exchanges between holdings. We calibrate the model using French epidemiological data. We then identify farm-level factors associated with the spreading potential of LA-MRSA in the network. We also show that, assuming control measures applied in a limited (n=100) number of farms, targeting farms depending on their centrality in the network is the only way to significantly reduce LA-MRSA global prevalence. Finally, we investigate the scenario of emergence of a new LA-MRSA strain, and find that the farms with the highest indegree would be the best sentinels for a targeted surveillance of such a strain’s introduction.

## 1. Introduction

Livestock farms are often at high epidemiological risk, in particular farms with a dense animal population ^1^. Disease spread control in livestock populations is a challenge for animal health and welfare ^2^, as well as for farmers in terms of economic and livelihood loss due to productivity drops ^3^. It may also be a concern for human health in the case of zoonotic diseases ^4^. In this context, over recent years, the worldwide spread of antimicrobial-resistant bacteria among livestock has emerged as a major threat that needs to be accounted for to fully understand the global increase of antimicrobial resistance in a one-health perspective ^5–7^.

Livestock-associated strains of methicillin-resistant *Staphylococcus aureus* (LA-MRSA) have been identified since the 2000’s in farm animals, especially pigs and veal calves, as well as in humans in occupational contact with livestock, in several European countries ^8–11^. *S. aureus* is a Gram-positive bacterium carried asymptomatically by a large portion of human populations ^12^, and is a frequent cause of opportunistic infections ^13^. Methicillin-resistant *S. aureus* (MRSA), that are multi-resistant to antibiotics including most β-lactams, were originally found in human populations in hospitals as sources of nosocomial infections, and then in the human community ^14^. LA-MRSA strains, mostly belonging to the ST398 subtype in Europe ^11^, were found able to spread to human populations not in direct contact with farm animals ^15–17^. As pigs are suspected to act as a reservoir for LA-MRSA ^18^, understanding the spread dynamics of LA-MRSA within and between pig farms is key to being able to design proper surveillance and control measures. In particular, the major impact of pig movements on the spread dynamics of LA-MRSA between farms has been underlined ^19–21^.

Over the last decade, a few mathematical models have been proposed to study the spread of LA-MRSA, within a single pig herd ^22,23^, at the scale of a pig farm network ^24–26^ and from a single pig farm to humans^27^. These models allowed to explore the implementation of control interventions based on reduced antimicrobial use, reduced mixing of pigs within and between farms, improved biosecurity, movements restrictions, as well as voluntary eradication of pigs. They estimated the risk of transmission to human populations even beyond farm communities if such farm-level control measures were not implemented. Finally, they showed that a description of LA-MRSA spread at both the within- and between-herd scales is necessary when assessing control strategies ^18,25^.

Here, we propose a novel model of MRSA spread in pigs, that combines within- and between-herds epidemiological and demographic dynamics. This model is based on data from a real pig farms network at a country level and is applied to the French situation. Using the model, we identify farm-related factors associated with extensive spread of LA-MRSA over the network. We next assess the effect of farm-level control measures on the network spread of LA-MRSA. Finally we optimize sentinel farms selection when designing active surveillance of a new strain in the network.

In the remainder of the text, the term “MRSA” will always refer to LA-MRSA. The term “sows” will always refer to the breeding sows aimed at farrowing. The term “gilt” will refer to young sows destined to replace older sows for farrowing.

## 2. Methods

### 2.1. Definitions

Several types of pig farms are defined, depending on the purpose of the farms (breeding or production) and the rearing stage at which they take pigs over (Fig 1). Breeding farms, including nucleus (SEL) and multiplier (MU) farms, are defined as farms exporting breeding sows (28-week old gilts). Other farms, including farrowing (FA), farrowing-post-weaning (FPW), post-weaning (PW), post-weaning-finishing (PWF), finishing (FI) and farrowing-to-finishing (FF) farms, are production farms and aim at producing pigs that will be slaughtered when they are 28 weeks old. Breeding farms include a breeding herd composed of sows aimed at producing new piglets. This is also the case for some production farms: types FA, FPW and FF. The fattening herd is composed of pigs raised to be sent to slaughterhouse, or, in the case of breeding farms, exported to other farms for renewing their breeding herd (see Supplementary Material S2).

**Fig 1.**
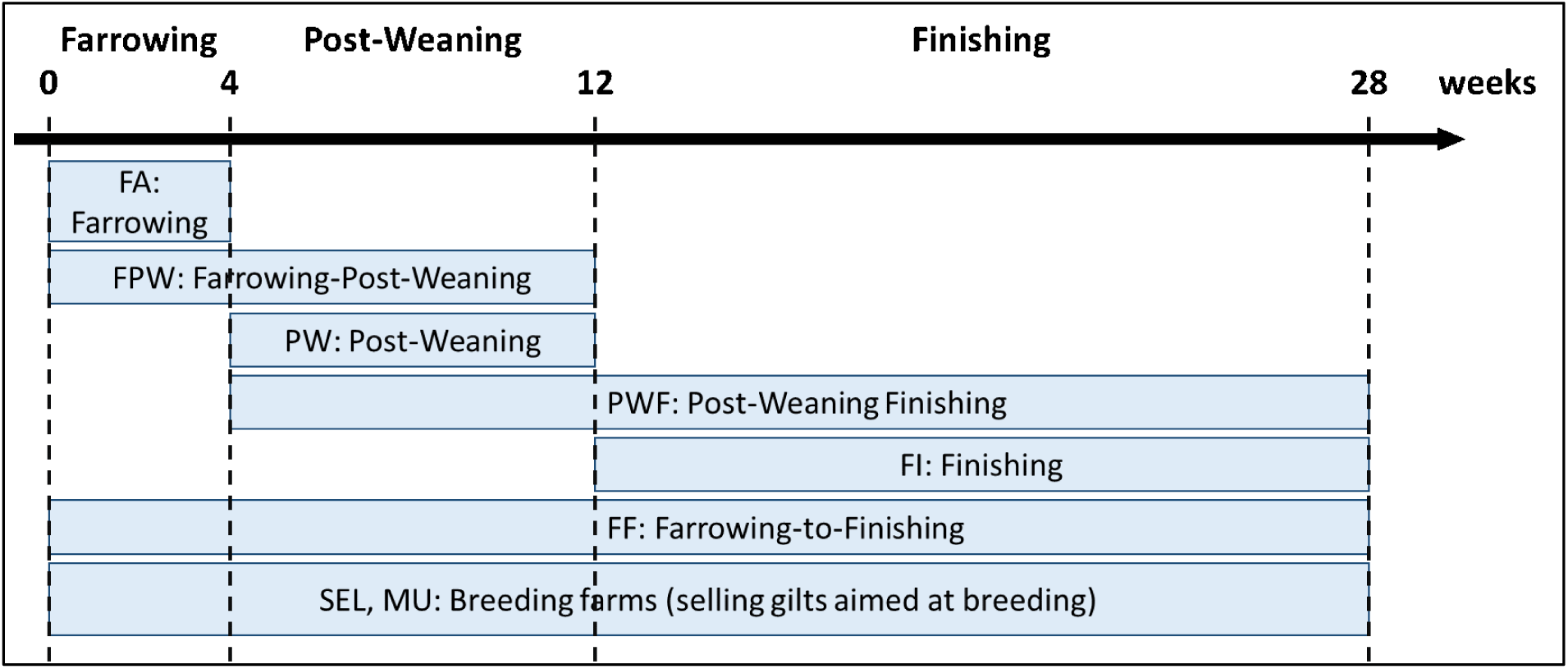
Types of pig farms. The pig production steps each farm type assumes are detailed.

### 2.2. Data on the French pig sector

The National Swine Identification Database (BDPORC) has recorded pig movements in France since 2010, allowing to reconstruct the network of French pig farms, which was described previously ^28^. This network was shown to be stable over time, with similar active nodes, network properties and connected components ^28^. Here, we used time-aggregated data from BDPORC for the full year 2014: the characteristics of all French pig farms, including their type of activity and size (Fig 2a); and all pig movements reported at the batch level (Fig 2b). Because we aimed to study the spread of MRSA between farms through pig movements only, the database was screened and filtered to exclude types of holdings for which pig exportations to other farms were very slight or absent (small farms, boar stations, wild boar farms, trade operators and slaughterhouses) ^28^, and types of movements that were rare and not consistent with farm types (e.g. fattening pig movements from finishing to multiplier farms). This resulted in 10,542 farms, which is consistent with the number of active nodes in the French pig industry identified in ^28^.

**Fig 2.**
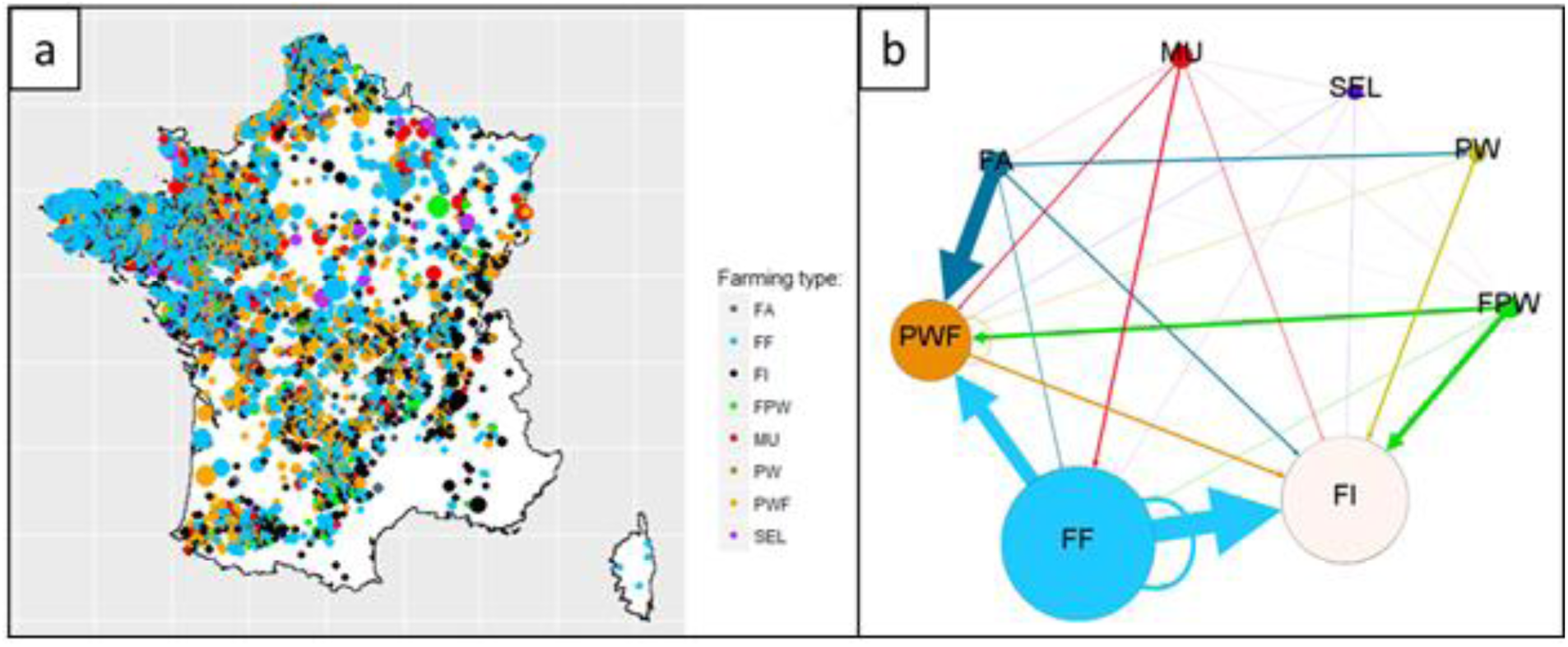
Characteristics of the French pig farm network. a) Map of French pig farms in 2014. Dot color represents the farm type (FA Farrowing, FF Farrowing-to-Finishing, FI Finishing, FPW Farrowing-Post-Weaning, MU Multiplier, PW Post-Weaning, PWF Post-Weaning Finishing, SEL Nucleus), and dot size is proportional to the number of pigs on site. b) Weighted oriented network of pig exchanges between farm types. Node size is proportional to the number of farms of the corresponding type; arrow size is proportional to and the number of pig moves between the two farm types.

### 2.3. Network analysis

In our analyses, we calculated several centrality indicators for farms within the network: outdegree, indegree, outflux, influx, betweenness, closeness, coreness and eigenvector centrality. The outdegree of farm A is the number of farms to which farm A exports pigs. The indegree of farm A is the number of farms from which farm A imports pigs. The outflux is defined as the number of pigs exported by farm A to other farms. The betweenness is the number of directed geodesics – i.e. directed shortest paths between each pair of nodes – going through farm A. Definitions for all these indicators are provided in the Supplementary Material S1.

### 2.4. Dynamic model description

We built a discrete dynamic stochastic model of the spread of MRSA within pig farms and between them over a connected network. Two distinct processes were modelled: MRSA colonization and transmission dynamics on the one hand, and pig demographics on the second hand. Both were modelled within-farm and at the network scale. Model parameters are summarized in Table 1, along with their assumed values.

**Table 1.** Model parameters and their assumed values.

| Variable | Description | Value | Source |
| --- | --- | --- | --- |
| $N_f$ | Number of farms in the network | 10,542 | BDPORC |
| $N_b$ | Number of farms in the network housing breeding sows | 5,102 | BDPORC |
| $D_{prod}$ | Duration of a full production cycle | 28 weeks | <sup>29</sup> |
| $\beta$ | Transmission rate of LA-MRSA between 2 pigs | 0.26 week<sup>-1</sup> | Calibrated |
| $d$ | Average LA-MRSA colonization duration in pigs | 4.3 week | <sup>30</sup> |
| $C_{STP}$ | Sow-to-piglets contamination probability | 1 | <sup>31</sup> |
| $P_{exp}$ | Proportion of fattening pigs exported at the end of each production step<br>(4 weeks or 12 weeks of age) | farm-specific | BDPORC |
| $P_{rep}$ | Proportion of sows replaced by gilts every 4 weeks | 3.19% | <sup>29</sup> |
| $P_{farms}$ | Proportion of farms contaminated by MRSA at $t=0$ | 5% | <sup>32,33</sup> |
| $P_{col farms}$ | Proportion of pigs colonized by MRSA in contaminated farms at $t=0$ | 16% | <sup>32,33</sup> |
| $P_{col total}$ | Global proportion of pigs colonized by MRSA at $t=0$ | 5% * 16% = 0.8% | |

#### 2.4.1. Demographic model

Demographic processes were modelled deterministically, at both the within-farm and between-farm levels.

The model formalized both breeding and fattening herds within farms (Fig 3a and Supplementary Material S2). The within-farm structure divided farms into 4 sectors: the gestation sector (GS), farrowing sector (FAS), post-weaning sector (PWS) and finishing sector (FIS). GS and FAS housed breeding sows. FAS, PWS and FIS housed fattening pigs of increasing age. Sectors were divided again into rooms: 4 rooms in GS, 1 in FAS, 2 in PWS and 4 in FIS (Fig 3a).

**Fig 3.**
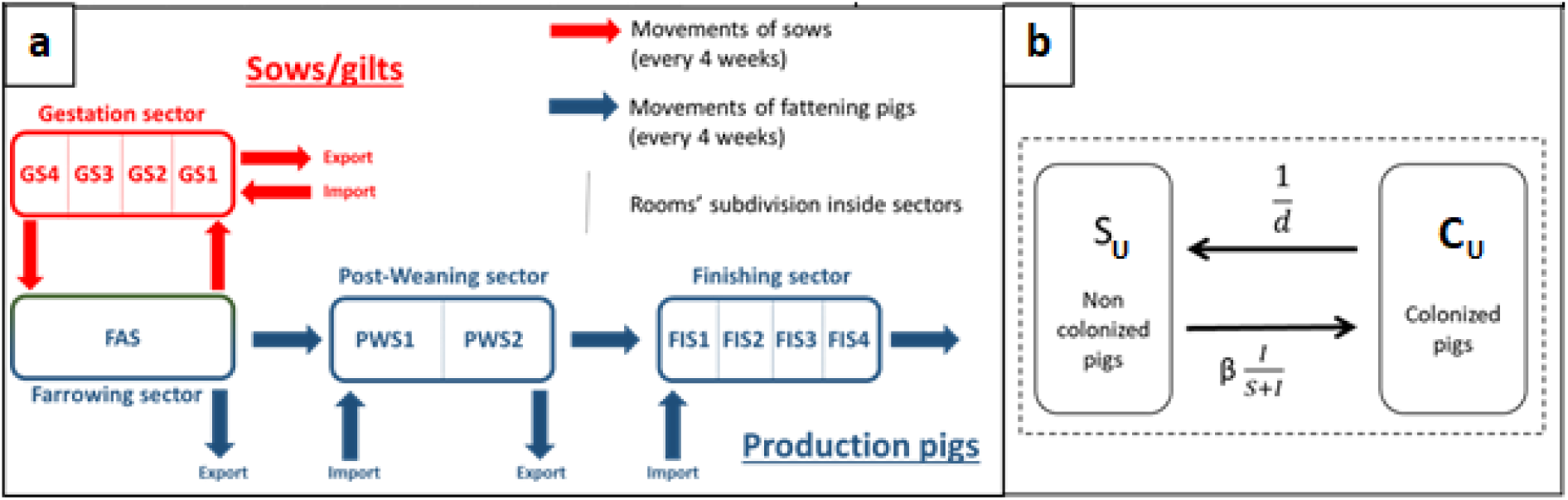
Mathematical model components. a) Demographic model at the within-farm scale. Each farm is divided into 4 sectors (GS, FAS, PWS, FIS), divided again into rooms (resp. 4, 1, 2 and 4). b) Epidemiological model within a given farm sector. S_U_ (resp. C_U_) is the number of non-colonized (resp. colonized) pigs in the sector.

We modelled a 7-batch rearing system, the most common pig management system in France ^34^, based on a 4-week cycle for pig movements. Each room in the FAS, PWS and FIS sectors housed a single batch. Therefore, piglets stayed 4 weeks in the FAS with their mother, then 8 weeks in the PWS, and finally 16 weeks in the FIS. In total, the production cycle for fattening pigs was assumed to last for 28 weeks (Fig 1), and farm size was supposed to be constant over time.

Pig movements between farms directly replayed the pig transfer network described above. Two types of pig exchanges between farms were possible every 4 weeks. On the one hand, gilts were transferred from breeding farms to replace a proportion *P*_*rep*_ of breeding sows in production farms. *P*_*rep*_ was assumed to be on average equal to 3.19%, based on 2014 French data in which 41.5% of sows in a sow herd were shown to be replaced over the entire year ^29^. Self-renewal of sows was also possible: in this case, the replacing gilts were randomly chosen among 28-week old pigs within their own farm’s finishing sector (FIS4 in Fig 3a). On the second hand, batches of fattening pigs could be moved between farms at two ages corresponding to two production steps: 4 weeks old (right after weaning), or 12 weeks old (for finishing). The proportion *P*_*exp,farm*_ of exported pigs from a given batch depended on the farm category. For instance, 100% of 4-week old piglets were exported from Farrowing farms, because these farms do not raise older pigs. Conversely, Farrowing-to-Finishing farms were assumed to export 12-week old pigs only. For each farm and at each production step, the number of pigs exported was computed from the network data and *P*_*exp, farm*_. When the exportation proportion could not directly be set to 0% or 100%, it was computed as follows based on the BDPORC data:

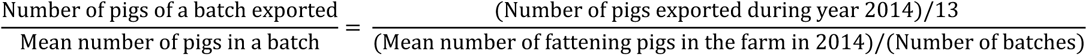

The 4-week periodicity of pig exchanges was systematically shifted among farms, so that they did not occur at the same time in all farms.

The resulting contact matrices between farm categories are consistent with data published by the French pig industry on piglets’ movements between categories of farms ^35^. More details on the between-farms model, including contact matrices between farm categories, are available from the Supplementary Material S2.

#### 2.4.2. Epidemiological model

Pig-to-pig MRSA transmission was modelled at the within-farm scale, through a discrete time stochastic Susceptible-Colonized-Susceptible (SCS) model (Fig 3b). Transmission could only occur within the same farm sector. Thus, the only way for MRSA to spread between farms was through movements of colonized pigs. Pigs were assumed not to change MRSA status during their between-farm transportation.

For any week *t*, the susceptible (i.e. non-colonized) and colonized pig populations within a given farm sector *U* at week *t+1* were calculated as follows:

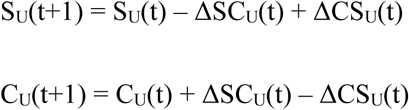

where S_U_(t) is the number of susceptible pigs in sector *U* at week *t*, and C_U_(t) the number of colonized pigs in sector *U* at week *t*. ΔSC_U_(t) and ΔCS_U_(t) represent respectively the flux from S_U_ to C_U_ state, and from C_U_ to S_U_ state, at week *t*. They were drawn from binomial distributions as follows:

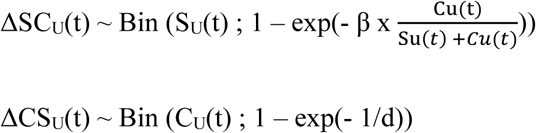

where β is the MRSA transmission rate between pigs of the same sector (assumed constant across all ages and all farms), and d is the average duration of MRSA colonization of a pig. In previous experimental work ^30^, MRSA was shown to persist in half of the inoculated pigs for at least 30 days (d = 4.3 weeks).

In farrowing sectors, because of close proximity between a sow and its piglets, we assumed that piglets had the same MRSA status (colonized or susceptible) as their mother, based on published experimental data ^31^. This status was assumed to be maintained during the entire farrowing step in both the sow and its piglets, due to constant recontamination occurring between them.

### 2.5. Model simulations

All simulations and analyses were performed using R version 3.4.3 “Kite-Eating Tree” and packages *mc2d, parallel, igraph, Hmisc* and *mratios*. Simulations were generated using discrete time steps of one week.

We first initialized the network as MRSA-free and ran the model until the pig herd populations stabilized. We then studied the spread of MRSA over the network using two distinct scenarios.

#### 2.5.1. The “realistic scenario”

In scenario 1, the “realistic scenario”, we used data from a cross-sectional study led by the French agency for food safety in 2007 to assess MRSA carriage in pigs in France ^32,33^. In this study, 5% of pig farms were found to be MRSA-positive (i.e. housing at least one MRSA-positive pig), and 0.8% of sampled pigs were MRSA-positive. To simulate this situation, at *t*=0, we randomly selected 5% of the farms to be MRSA-positive. In these farms, we set 16% of randomly selected pigs of all ages as MRSA colonized, to result in 0.8% of total prevalence in pigs.

Scenario 1 was used for calibrating β, and for assessing the impact of targeted control measures in an endemic situation. In this scenario, due to stochasticity, 300 repetitions of the model were simulated, as this value was found to be enough to hold stable the mean and variance of model outputs, namely the percentage of farms contaminated (PFC), the total prevalence in pigs and the prevalence of pigs heading for slaughterhouse.

#### 2.5.2. The “introduction scenario”

In scenario 2, the “introduction scenario”, we assumed an initial MRSA-free situation and simulated the introduction of a single MRSA-positive group of gilts in a single given farm (the seed). At *t*=0, we set 3.19% (that is *P*_*rep*_, the portion of sows replaced by gilts every 4 weeks in our model, see Table 1) of the breeding herd of the seed farm as MRSA-colonized, among sows entering gestation. This MRSA introduction process was repeated in simulations starting from each of the 5,102 farms housing breeding sows.

Scenario 2 was used for identifying factors associated with the spreading potential of farms, and for selecting efficient sentinel farms to detect the dissemination of a new MRSA strain over the pig farm network. In this scenario, for each seed farm, 50 stochastic repetitions were simulated, this number being sufficient to hold model outputs’ mean and variance stable.

### 2.6. Model and data analysis

#### 2.6.1. Model calibration

We calibrated the value of the transmission rate, β using scenario 1. We aimed to select the value of β minimizing residuals between the mean predicted prevalence in pigs of all farm sectors at steady state (among all simulations), and the reported carriage of 0.8% ^32,33^.

#### 2.6.2. Analysing the spreading potential of seed farms

We performed a multivariate analysis to assess the relationship between the spreading potential of the seed farm in scenario 2. Spreading potential was measured through the percentage of farms contaminated (PFC) at *t*=52 weeks (one year) predicted with this scenario, and the seed farm’s characteristics. Investigated characteristics were the seed’s breeding herd size, production herd size, farming category (breeding farm, i.e. Multiplier or Nucleus, VS production farm), and 8 network centrality indicators: outdegree, indegree, betweenness, eigenvector centrality, closeness, coreness, outflux and influx.

The variables associated with the PFC in univariate linear regression with a p-value<0.2 were selected for the multivariate analysis. Variable selection in the multivariate linear models was performed using a stepwise selection procedure algorithm, using Akaike’s information criterion (AIC). We checked the multicollinearity among explanatory variables selected in the multivariate analysis by computing the Variance Inflation Factors (VIF). In case variables showed a VIF > 10, they were excluded from the multivariate model, and the stepwise algorithm and VIF checking procedure was repeated without them.

For illustrative purposes, we focused more specifically on the association between PFC and the seed farm’s outdegree and category.

#### 2.6.3. Assessing the impact of targeted control measures

Assuming that efficient within-farm control measures to reduce MRSA spread exist, one may need to identify which farms should be targeted preferentially, through a program supporting specific farmers to implement them for instance. In this part, we used our model to determine the types of farms in which these measures should be selectively applied to enhance their effectiveness at the national production level. We compared the impact of farm-level control measures targeting 100 farms with either the highest indegree, the highest outdegree, the highest outflux or the highest betweenness, with that of the same control measures implemented in 100 random farms. As in ^23^, we assumed that control measures reduced the transmission parameter β in farms. In the “realistic” scenario 1, once the prevalence had plateaued, at *t*=208 weeks, we reduced the transmission parameter in the 100 selected farms (targeted or randomly selected). Two levels of reduction were assessed: 25% (leading to β = 0.19/week) and 50% (leading to β = 0.13/week), and compared to the “no action” baseline value β = 0.26/week. We investigated the total prevalence of MRSA colonization in pigs of all ages and the PFC at different times after the introduction of the control measures.

#### 2.6.4. Sentinel selection for targeted surveillance

We used our model to seek the best method to select sentinel farms to perform targeted surveillance of incursions of a new MRSA strain in the network, based on “introduction” scenario 2 simulating outbreaks starting from all farms housing sows. We ranked farms in priority lists: for instance, if for economic or practical reasons, only N farms can be monitored regularly, which should be the first N farms to be monitored to obtain the most efficient surveillance? We investigated three criteria for surveillance efficiency: the percentage of MRSA incursions detected, the time before MRSA detection, and the PFC at detection (that is, the outbreak size at detection). As in ^36^, we calculated the Kendall correlation between these 3 criteria. We compared 10 distinct methods of targeted sentinel selection ^36–39^ to a random selection of sentinels. The first 8 of these methods were simply based on sorting farms by decreasing values for network centrality measures (see Supplementary Material S1 for definitions): outdegree, indegree, betweenness, eigenvector centrality, closeness, coreness, outflux and influx ^36,38^. The 9^th^ method was the “Invasion paths” method, described in ^37^ and ^39^. The principle of this method is to cluster farms based on the similarity of the paths that would potentially take pathogens in the oriented network (details of the method are provided in Supplementary Material S3). In the last method, we set a priority list by selecting farms that maximized alternatively the following criteria: 1) the best farm in term of percentage of detected incursions, 2) the best in term of time before detection, 3) the 2^nd^ best in term of percentage of detected incursions, etc. We called this the “Alternated method”. For all these methods, we assessed the surveillance efficiency depending on three numbers of sentinels monitored: 30, 60 and 120 sentinel farms.

## 3. Results

### 3.1. Model calibration and baseline predictions

Under “realistic” scenario 1, for values of β <0.3/week, a non-null steady-state for the PFC and the national prevalence in pigs could not be reached, due to local extinctions in farms (see Supplementary Material S4). Irrespective of the value of β, the prevalence plateaued during the 4th year of simulation, before a slow and steady decrease (see Supplementary Material S4). Hence, we selected the value minimizing residuals between the predicted prevalence in pigs of all farm sectors, and the observed carriage of 0.8%, between *t*=156 weeks and *t*=208 weeks (i.e. in the 4th year of simulation). The calibrated baseline value of the transmission parameter is β=0.26/week. Simulations show that with this transmission parameter value, the predicted prevalence of MRSA in pigs in scenario 1 remains close to 0.8% for more than 8 years of simulations (see Supplementary Material S4).

Model simulations showed high stochasticity at the farm level, with frequent extinctions and transmissions. As an illustration, Figure 4 depicts the predicted prevalence of MRSA colonization among the pigs from 4 selected farms in our model under the “introduction” scenario 2, as a function of time (Fig 4a), as well as pig movements between these same farms (Fig 4b).

**Fig 4.**
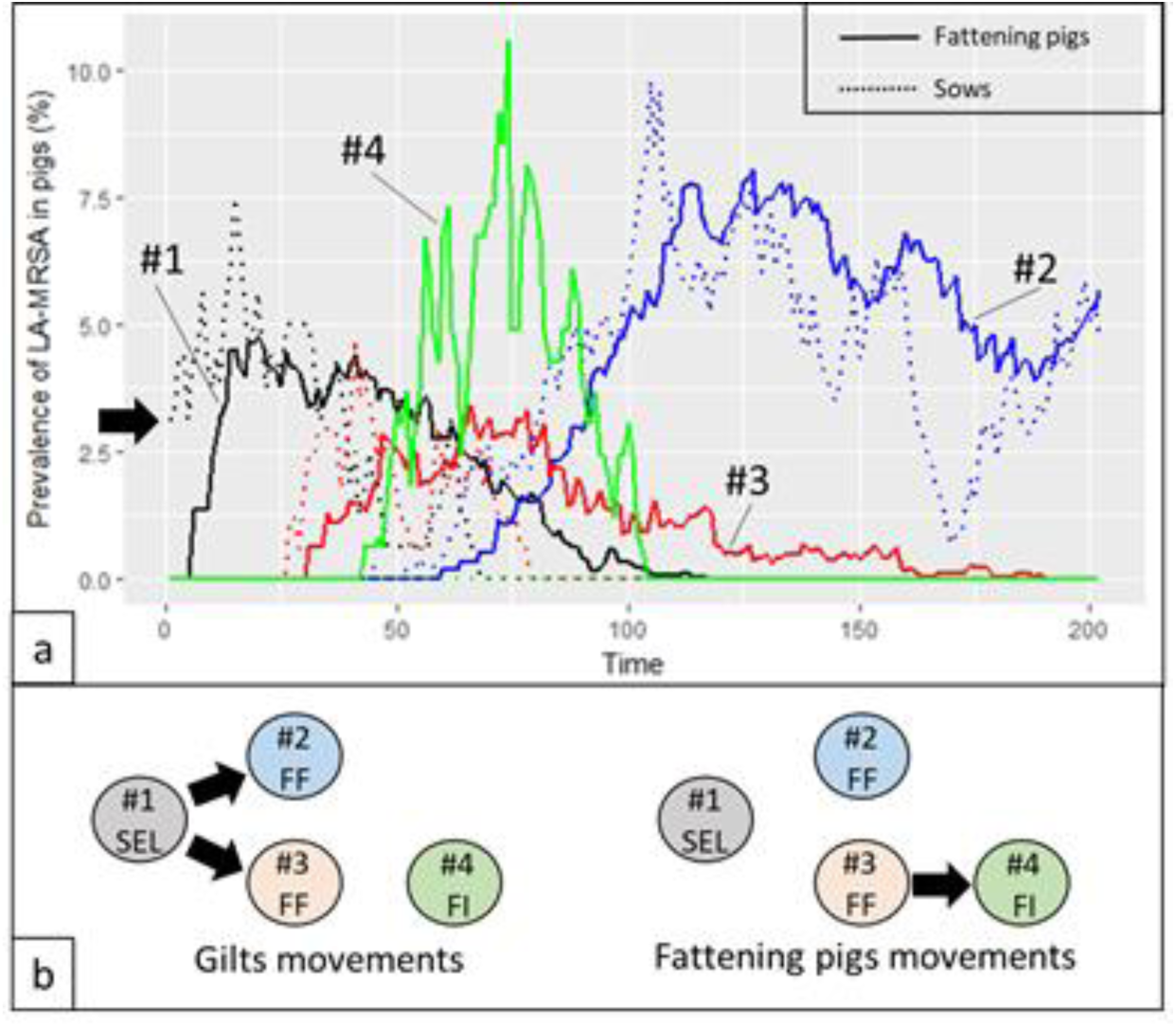
Illustration of model simulations. MRSA spread in 4 farms in our model based on a single stochastic simulation: a nucleus farm (SEL), where MRSA was seeded, and three production farms (2 farrowing-to-finishing farms (FF) and a finishing farm (FI)). a) Time changes in MRSA prevalence in the breeding (dotted line) and fattening (solid line) herd of each farm. b) Pig movements between the 4 farms, according to the database, differentiating gilts and fattening pig movements.

Farm #1 (a Nucleus farm) was the seed farm in this simulation: 3.19% of its breeding herd was contaminated at initialization. Farms #2 and 3 (Farrowing-to-Finishing farms) were first contaminated through importation of colonized gilts. Then, MRSA reached their fattening herd. Farm #4 (a finishing farm) was however contaminated through importation of fattening pigs. Due to stochasticity, extinction was observed in some farms (see Supplementary Material S4).

### 3.2. Factors associated with the spreading potential of seed farms

Results of the multivariate analysis assessing the impact of the characteristics of the seed farm on outbreak size (measured through the predicted PFC after 1 year) are detailed in Supplementary Material S5. Being a breeding farm (as opposed to a production farm), as well as higher values of production herd size, outdegree, betweenness, closeness, coreness and outflux, were significantly associated with a higher spreading potential (p-values < 10^−12^). A higher value of indegree was significantly associated with a lower spreading potential (p-value < 10^−15^). In the final multivariate linear model, all VIF were < 2.5, showing a limited multicollinearity among explanatory variables.

As an illustration, Figure 5 highlights the effect of the seed’s farming category and outdegree on two indicators after 1 year: the PFC and the prevalence of MRSA in pigs heading for slaughterhouse, just before transportation. Irrespective of the seed farm, the outbreak size in this scenario remained limited: the average proportion of farms contaminated after one year was 0.03% (SD: 0.05) (n=3.3 farms), and the maximum was 0.62% (n=65.3 farms). Moreover, after 1 year, the average percentage of pigs colonized by MRSA sent to slaughterhouse was 9.6×10^−4^ % per week (SD: 1.0×10^−3^), and the maximum was 0.01% per week (Fig 5).

**Fig 5.**
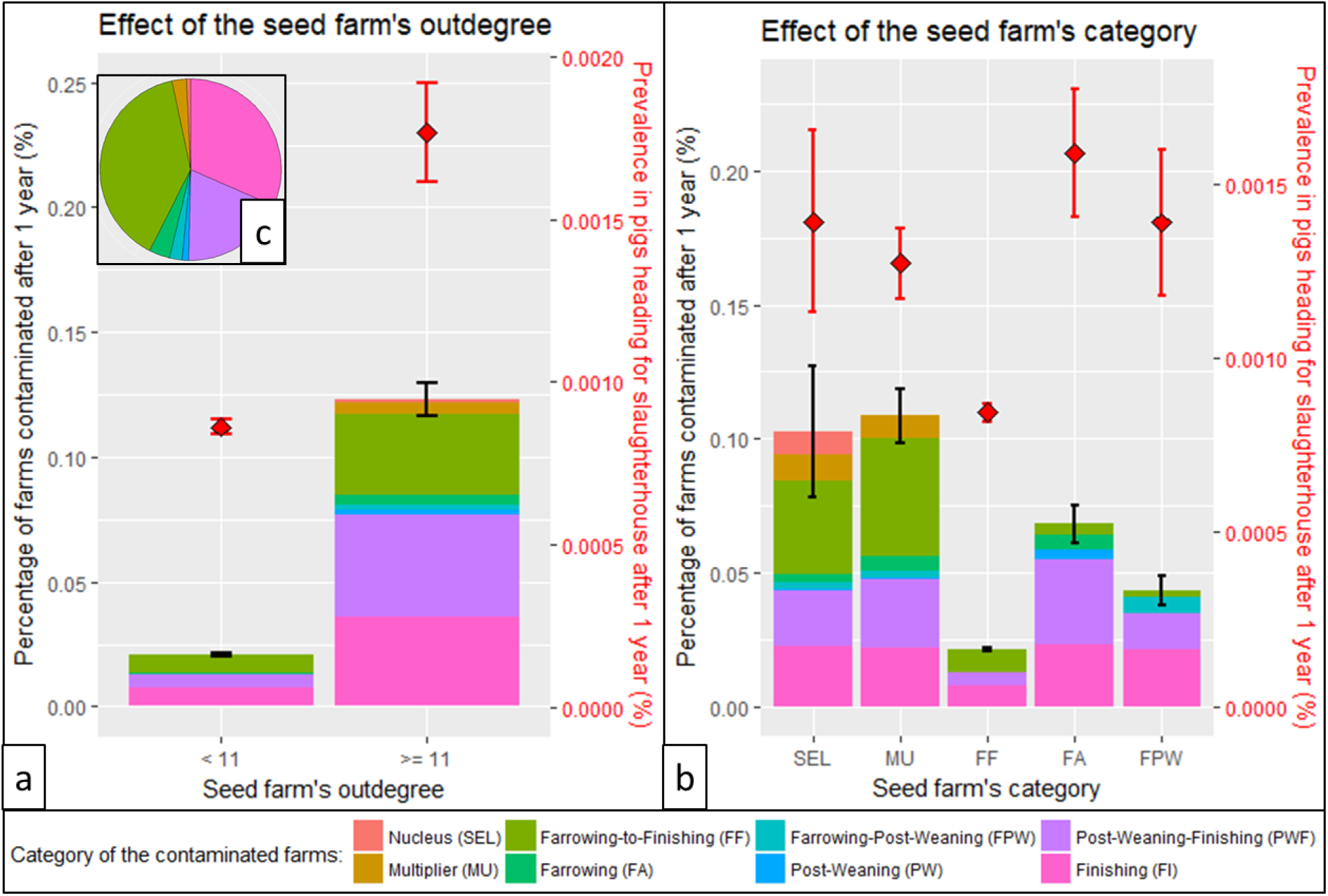
Impact of seed farm characteristics on its spreading potential. Using the “introduction” scenario 2, the effect of the seed farm’s outdegree (a), and category (b), on two outputs after 1 year is depicted: the proportion of farms contaminated (PFC, left axis, bar graphs) and the national prevalence of MRSA in pigs heading for slaughterhouse (right axis, red diamonds dots). We also show in which proportions the different categories of farms (including the seed farm) are contaminated. Intervals are 95% confidence intervals of the one-sample t-test on the mean of the output (PFC and prevalence at slaughter age) for farms of a given category or outdegree.

Regarding the effect of the seed’s outdegree, we compared the spreading potential of the 10% of seed farms with the highest outdegree (category “>= 11”) with that of other seed farms (category “< 11”) (Fig 5a). The PFC predicted 1 year after MRSA introduction significantly increased with the seed farm’s outdegree (Welch t-tests between categories “< 11” and “>= 11”, p-value<10^−15^). It also depended on the seed farm’s category, with higher PFC predicted when MRSA was seeded in breeding farms (with no statistical difference between nucleus and multiplier farms), and lower PFC when MRSA was seeded in production farms (Welch t-test, p<10^−15^) (Fig 5b). Among production farms, MRSA introduction in a farrowing farm led to a higher PFC than introduction in a farrowing-post-weaning farm (Welch t-test, p<10^−7^), and the lowest PFC was predicted when the seed farm was a farrowing-to-finishing farm (Welch t-tests between categories “FF”, and “FA” or “FPW”, both p<10^−13^).

1 year after MRSA introduction, the predicted prevalence among pigs at slaughter age also significantly increased with the seed farm’s outdegree (Welch t-tests between categories “<11” and “>= 11”, p-value<10^−15^) (Fig 5a). It was significantly lower when the seed farm was a farrowing-to-finishing farm (Welch t-tests between categories “FF”, and all others, all p<10^−4^). Introducing MRSA in a Multiplier farm led to a higher prevalence among pigs heading to slaughterhouse than in a Farrowing farm (Welch t-test, p<0.01) (Fig 5b).

The proportion of each farm category within the contaminated farms depended on seed farm’s characteristics, and could highly vary from the proportions of farm categories in the network (Fig 5). In particular, seeding MRSA in a Nucleus farm led to a proportion of Multiplier farms among contaminated farms about 4 times higher than the proportion of Multiplier farms in the network (10.0% against 2.6%). On the contrary, whereas Farrowing-to-Finishing farms represented 39.2% of farms in the network, they were only 5.6% – i.e. a 7-fold decrease – among contaminated farms when the seed farm was a Farrowing farm.

### 3.3. Impact of targeting farms when implementing control measures

Based on “realistic” scenario 1, Figure 6 compares the impact of applying farm-level control measures – i.e. reducing the transmission parameter β – in 100 farms, either chosen randomly, or targeting the ones with the highest outdegree. Irrespective of the assumed intensity of control measures (amount of β reduction), the delay after their implementation and the model output considered, control measures implemented in random farms had no significant impact on MRSA spread (ratio t-test comparing model predictions with and without control measures, p>0.05). On the contrary, a 25% (resp. 50%) β reduction in targeted farms significantly decreased MRSA prevalence, as soon as two (resp. one) years after the implementation of control measures (p<0.01) (Fig 6a). In this case, the impact of control measures increased with time passed since their implementation. For instance, a 25% reduction of within-farm MRSA transmission in targeted farms led to an 8.7% (Fiellers 95% confidence interval (CI) [3.1%; 14.0%]) reduction in the predicted overall MRSA prevalence 2 years after control measures implementation, and to an 18.1% (Fiellers 95% CI [12.6%; 23.4%]) reduction after 4 years, compared to the “no action” baseline case. Similar results were obtained regarding the PFC (Fig 6b). Comparable results were also obtained when considering the 100 farms with the highest betweenness or the 100 farms with the highest outflux as targets for the implementation of within-farm control measures (Supplementary Material S6).

**Fig 6.**
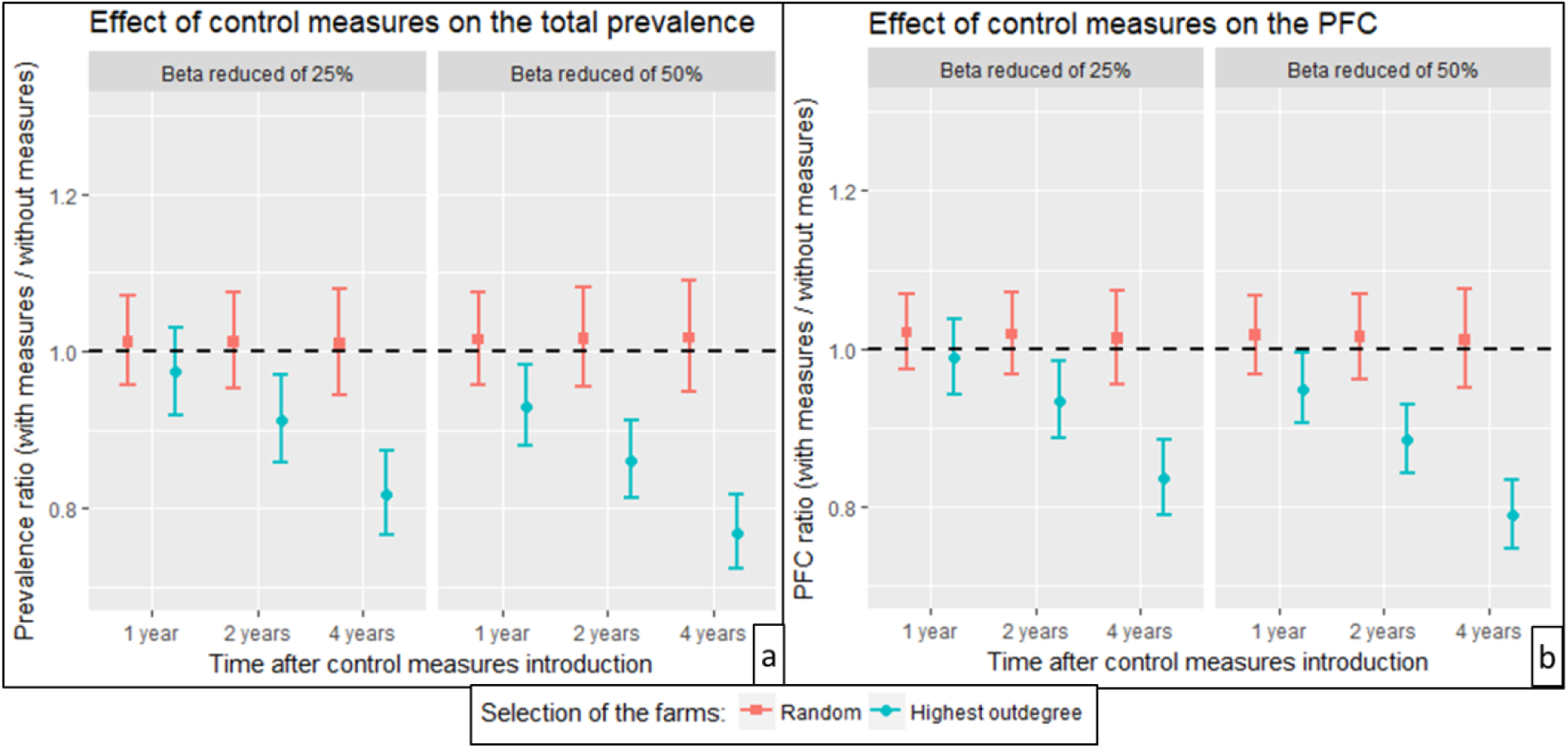
Impact of targeted control measures. Impact of control measures implemented in 100 farms selected at random (red bars) or in the 100 farms with the highest outdegree (blue bars). The ratio “Output value with control measures / Output value without control measures” is depicted at different times (1, 2 and 4 years after control measures implementation) for two model outputs: (a) Total prevalence in pigs and (b) Percentage of Farms Contaminated (PFC). “Realistic” scenario 1 was simulated. Two levels of control measure intensity are considered: reducing the transmission parameter β by 25% or 50%. Intervals are Fiellers 95% confidence intervals of the t-test for the ratio of two means: the mean values of outputs of 300 model iterations for the case control measures are applied VS the case control measures are not applied.

### 3.4. Impact of sentinel farm selection for targeted surveillance

Figure 7 compares the performance of several methods for sentinel farm selection, in terms of three criteria for surveillance efficiency: the percentage of detected incursions (Fig 7a), to be maximized, and the time before detection (Fig 7b) and the PFC at detection (Fig 7c), to be minimized.

**Fig 7.**
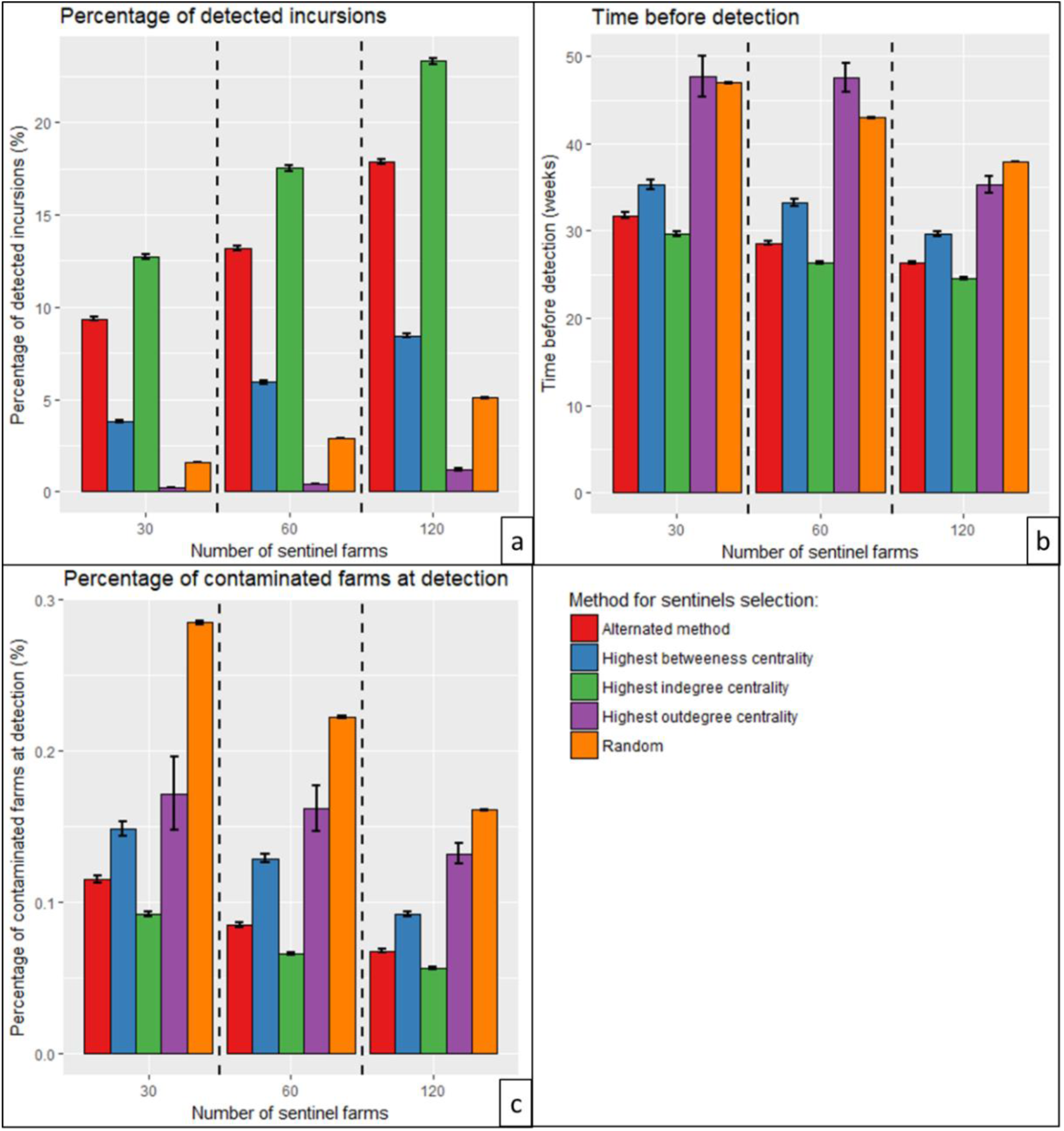
Comparison of the efficiency of several methods for sentinel selection. Three efficiency criteria are depicted: Percentage of MRSA incursions detected (a), Time before detection (b), and Percentage of farms contaminated (PFC) at detection (c). Several sizes of sentinel sets were tested: 30, 60 and 120 sentinel farms. For each set of sentinel farms, the surveillance performance was assessed based on the average of model simulations of MRSA introduction starting from all farms housing breeding sows in the network (scenario 2). Intervals are 95% confidence intervals of the t-test. 100 different sets of sentinels were considered with the “Random” choice method.

Irrespective of the selection method, all three efficiency criteria improved when the number of sentinel farms increased. For instance, with the highest indegree method, a 4-fold increase in the number of sentinels (from 30 to 120) multiplied the percentage of detected incursions by 1.8 (from 12.7% to 23.3%), and allowed to detect the incursion 5.1 weeks earlier (from 29.7 to 24.6 weeks).

For all efficiency criteria, the best method for sentinel selection was the one based on the highest indegree, and the second best was the “Alternated method”. For instance, monitoring the 120 farms with the highest indegree allowed to detect an incursion after 24.6 weeks on average (95% prediction interval [10.0; 73.0]), 13.4 weeks earlier than when sentinels were chosen at random; upon detection, the average PFC was 0.06% (95% prediction interval [0.01%; 0.28%]) (n=6.0 farms), 2.8-fold less than when sentinels were chosen at random. Other sentinel selection methods that proved less performant than the highest indegree on all 3 criteria are displayed in Supplementary Material S7.

## 4. Discussion

Pig movements between farms are believed to be one of the main factors of MRSA dynamics in the pig industry ^21^. Moreover, recent modelling work underlined the importance of considering within-farm dynamics when studying between-farms spread of MRSA ^25^. In this study, we propose a mathematical model combining the within-farm and the between-farm levels to simulate the spread of MRSA in the French pig industry. We highlight the importance of individual and network-based farm characteristics on their role in the global MRSA epidemic. We compare several scenarios to target within-farm control measures implementation, and to identify pertinent sentinel farms for MRSA surveillance.

### 4.1. Main findings

The “introduction” scenario we proposed in our analyses, in which MRSA is introduced in France through a single seed farm, is not realistic, in the sense that it leads to a maximum PFC of 0.62% (n=65 farms), far from the 5% observed in France in 2007 ^32,33^. This reflects the fact that MRSA was probably introduced in French pig farms through several distinct farms, and on repeated occasions over time, probably as was shown in Denmark ^21^. However, this scenario is still useful for two main reasons. First, it allowed us to investigate the specific spreading potential of different farm types. Second, it may realistically simulate the situation where a hypothetical new MRSA strain should arrive in France, and propose criteria for sentinel surveillance of such incursions.

We found that MRSA spread in the pig industry largely depends on the characteristics of the farm in which it is first introduced, namely its size, category and its centrality in the network. Outbreaks originating from breeding farms (Multiplier or Nucleus) or from farms with high outdegree led to a significantly higher proportion of farms contaminated (PFC) than outbreaks originating from production farms (Farrowing, Farrowing-to-Finishing or Farrowing-Post-Weaning) and from farms with lower outdegree (Fig 5). Results regarding the prevalence in pigs heading for slaughterhouse are less clear. This may be due to the low predicted prevalence values in pigs at slaughter age.

One strategy to set up global infectious diseases surveillance over the network is to choose sentinel nodes that are closely monitored. The choice of these sentinel farms is however challenging as, depending on this choice, the reactivity in sending the alert can vary. We proposed and assessed several methods to help select sentinel farms that may be used in a surveillance system. Because disease surveillance in livestock meets limitations as only a sample of the total pig industry can be regularly monitored, for costs reasons, we simulated the monitoring of 30 to 120 sentinel farms only. There was a high Kendall’s correlation coefficient value between the 3 surveillance efficiency criteria we considered (Percentage of detected incursions VS Time before detection: 0.81 (p-value < 10^−15^); Percentage of detected incursions VS Number of contaminated farms at detection: 0.71 (p-value < 10^−15^); Time before detection VS Number of contaminated farms at detection: 0.82 (p-value < 10^−15^)). This justifies why we tested the “Alternated method” for sentinel selection. Indeed, these correlations can lead to difficulty in optimizing simultaneously the percentage of detected incursions, the time before detection and the outbreak size (PFC) at detection, as evidenced by earlier work ^36^. This alternated method of sentinel selection proved to be rather efficient (Fig 7). However, we found that the sentinel selection method based on choosing the farms with the highest indegree was even more efficient, irrespective of the chosen criteria and the number of sentinel farms monitored.

Finally, different types of control measures of MRSA spread within farms have been suggested ^23,40,41^. Earlier studies ^26^ have suggested that the implementation of within-farm control measures could have an effect on the global pig farms network. In the hypothesis where efficient within-farm control measures were available, we showed that targeting certain categories of farms may increase their effectiveness at the network level. For the same “cost”, that is, the same amount of control interventions implemented to decrease MRSA’s within-herd transmission (e.g. a hypothetical program to enhance biosecurity in farms ^23^), our results suggest that it is more efficient to target pig farms sending pigs to the highest number of other farms (highest outdegree), pig farms exporting the highest number of pigs (highest outflux), or pig farms with the highest betweenness, than random farms (Fig 6 and Supplementary Material S6). Consistently with previous findings ^24^, our simulations also confirm that MRSA would be hard to eradicate in the pig industry. Indeed, in our simulations, the obtained reduction in total prevalence after 4 years is only 23%, even when reducing transmission by 50% in 100 well-targeted farms (with the highest outdegree), and assuming no other MRSA introduction occurs meanwhile in the farms’ network.

Finally, it should be noted that, among the 100 farms with the highest outdegree, outflux and betweenness respectively, only 1, 1 and 5 farms respectively were also part of the 100 farms with the highest indegree. This suggests that the farms monitored to detect a hypothetical introduction should not be the same than the ones targeted for within-farm control measures implementation once detection has occurred.

### 4.2. Main study limitations

This study has several limitations.

First, our calibrated value of the transmission parameter β – 0.26/week – was 7 times lower than the value estimated in a previous study ^42^, which was 0.21/day to 0.42/day. However, in our model, β accounted for global transmission of MRSA among a whole pig sector, including indirect transmission through the environment, whereas ^42^ only accounted for direct contacts between pigs. This could partly explain this lower value, along with the fact β was calibrated to reproduce a low observed carriage in France. What is more, in another study performed by the same team ^43^, the parameter calculated for transmission from other pig pens in the absence of antibiotics was 0.039/day, which is close to our calibrated value of 0.26/week (i.e. 0.037/day). The effect of variations of β on the model’s predictions are shown, for the “realistic” model, in Supplementary Material S4. The local extinctions we observed in some farms were likely due to the low value of our transmission parameter β, together with our model’s stochasticity and its within-farm structure, leading to smaller populations in contact (within sectors). While observed data suggest that pig farm can remain MRSA-positive for long time periods, such spontaneous extinctions have also been documented ^44^.

We assumed that pig farm sectors were spatially distinct enough, and the biosecurity on farm strong enough, for transmission to occur within sectors only, and not between sectors. In future work, it may however be interesting to compare a within-farm and a within-sector transmission parameter using longitudinal experimental data, as in ^42^, at the sector level. Furthermore, we chose not to take into account age differences in the transmission parameter, due to the limited carriage data available that did not allow us to calibrate several values for the transmission parameters. Provided age-specific MRSA prevalence data becomes available, this might prove an interesting addition to the model.

Second, we chose to select a value of MRSA colonization duration – 30 days – based on biological data ^30^, although other values were found in the literature, such as 17.4 days in ^42^. In comparison, the colonization duration we used may have the effect of favouring the spread of MRSA between farm sectors and between farms, as pigs carrying MRSA for a longer time were more likely to be transferred to another farm or farm sector while colonized. This might compensate the low value of β we used.

Third, we assumed that piglets had the same colonization status as their mother, based on ^31^, even though other experimental results ^45^ suggested that the association between sow and piglets colonization status may not be so direct. Using a probabilistic approach on the piglets’ colonization status would have potentially decreased the number of animals entering the production cycle while colonized by MRSA, and therefore the prevalence and PFC.

### 4.3. Perspectives and veterinary public health implications

Our results show that increasing the number of sentinel farms improves the surveillance efficiency, but more in-depth cost-effectiveness analyses would be useful to investigate what would be the most cost-efficient number of sentinel farms, regarding surveillance costs and veterinary public health objectives. It would also be interesting to compare a surveillance led in farms, versus a surveillance led at the slaughterhouse level. The hypothetical within-farm control measures we discussed may also have different implementation costs depending on farm size or category. Besides, the impact of potential control measures and surveillance on farm productivity and daily work, as well as on animal health and welfare, should always be carefully assessed before their implementation.

Furthermore, our findings in terms of MRSA prevalence among pigs sent to slaughterhouses should not be interpreted directly in term of contamination by MRSA of pork products intended for human consumption. Indeed, from the moment pigs leave the farm to the consumer’s fork, many factors, during transport, in the slaughterhouse and during food processing, are susceptible to affect the bacterial load in pork ^46^. In the future, these limitations should be addressed in a detailed risk assessment study to evaluate the risks for human health ensuing from the pig MRSA epidemic.

### 4.4. Conclusions

In conclusion, we show here how, using a multi-scale model of MRSA spread in French pig farms, criteria may be proposed to select which farms could be used as sentinels in a surveillance system, as well as which farms to apply control measures in. More thorough cost-effectiveness analyses, accounting for actual economic costs related to different types of control or monitoring interventions, would be necessary in the French context, similar to what was done recently in Denmark ^47^. However, this work has the potential to help better understand MRSA spread in the French pig industry. What is more, the methodology we proposed could also be applied to other asymptomatic bacteria or viruses circulating in pigs or other farm animals networks.

## Acknowledgments

We acknowledge BDPORC for providing the data and the doctoral program “Frontières de l’Innovation en Recherche et Éducation”. JB was funded by the INCEPTION project (PIA/ANR-16-CONV-0005), and his work was supported by internal resources of Institut Pasteur, the French National Institute of Health and Medical Research (Inserm) and the University of Versailles Saint-Quentin-en-Yvelines (UVSQ). Funding was also received from the French Government “Investissement d’Avenir” program Laboratoire d’Excellence “Integrative Biology of Emerging Infectious Diseases”, grant ANR-10-LABX-62-IBEID). The funders had no role in study design, data collection and analysis, decision to publish, or preparation of the manuscript.

## Authors’ contribution

MA, CC, PG, LO and LT conceptualized the project. LO and LT supervised the project. MA prepared and provided the dataset. JB, LO and LT designed the model and performed the statistical analysis. JB developed the model. MA and CC provided expertise on the pig industry and advice on the model. All authors approved the latest version of this article.

## Supplementary Material

S1. Definitions of network centrality indicators used in the article

S2. Details on the model

S3. Details on the “Invasion paths” method to select sentinel farms

S4. Model behaviour under the “realistic” scenario 1

S5. Results of the multivariate analysis to identify factors associated with the spreading potential of seed farms under the “introduction” scenario 2

S6. Additional results – Impact of targeted control measures

S7. Additional results – Sentinel selection for targeted surveillance

