## Supplementary Material for "Dynamics of Livestock-Associated Methicillin Resistant *Staphylococcus aureus* in pig farms networks: insight from mathematical modeling and French data"

### S1. Definitions of network centrality indicators used in the article

Centrality indicators measure the position of a node (here, a farm) relatively to other nodes within a network. We used the *igraph* R package to calculate them [1].

In our case, the ***outdegree*** of a farm A is the number of farms to which farm A exports pigs. The ***indegree*** is the number of farms from which farm A imports pigs. The ***outflux*** is the number of pigs exported by farm A to other farms. The ***influx*** is the number of pigs imported by farm A from other farms. The ***betweenness*** is the number of directed geodesics – i.e. directed shortest paths between each pair of nodes – going through farm A.

Farm A's ***closeness*** is defined as the inverse of the average length of the shortest paths linking farm A to every other node in the network. We consider movements going from and to farm A. Here, the closeness considered between two nodes without connecting path is calculated using the total number of nodes in the network instead of the path length.

The ***coreness*** of a node is based on the notion of k-core. The ***k-core*** of a network is the maximum sub-network in which all nodes have at least a degree of k (where degree = outdegree + indegree). The coreness of farm A is the value of k such that farm A belongs to the k-core of the network, but not to the (k+1)-core.

Finally, ***eigenvector centrality values*** are calculated from the network adjacency matrix. A high eigenvector centrality value for farm A represents the fact farm A is connected to nodes that are themselves connected to many nodes.

**S2. Details on the model**

1) Pig farm network data

The network is static (time-aggregated for year 2014). Database filtering was led in order to exclude farms with no incoming nor outgoing recorded movement, and to include only farms with the following types of activity: nucleus SEL, multiplication MU, farrowing FA, farrowing-to-finishing FF, finishing FI, farrowing-post-weaning FPW, post-weaning PW, post-weaning-finishing PWF (Table S1). In particular, farms defined as small farms (less than 80 pigs on site) were not included in the analysis.

***Table S1.*** *Categories of farms in the network database and in our model.*

<table>
<tr><th></th><th>Abbreviation</th><th>Category of farm</th></tr>
<tr><td rowspan="6">Production farms</td><td>FA</td><td>Farrowing</td></tr>
<tr><td>FPW</td><td>Farrowing-Post-Weaning</td></tr>
<tr><td>PW</td><td>Post-Weaning</td></tr>
<tr><td>PWF</td><td>Post-Weaning-Finishing</td></tr>
<tr><td>FI</td><td>Finishing</td></tr>
<tr><td>FF</td><td>Farrowing-to-Finishing</td></tr>
<tr><td rowspan="2">Breeding farms</td><td>MU</td><td>Multiplier</td></tr>
<tr><td>SEL</td><td>Nucleus</td></tr>
</table>

2) Within-farm demographic process

The within-farm model structure is the same for every farm in the network and is based on batch management. In the model, each farm is divided into 4 sectors (Gestation GS, Farrowing FAS, Post-Weaning PWS, Finishing FIS), each of them divided into respectively 4, 1, 2 and 4 room housing one batch each. This formalizes a 7 batches management, the most common pig herd management system in France [2]. However, the farm's category determines which sections are included in our model. For instance, the Post-Weaning-Finishing farms do not include GS and FAS. Each farm's demographic dynamic is based on a 4 week cycle: changes of sector, piglets' farrowing, shipment to slaughterhouse and sows' replacement occur every 4 weeks. A batch spends 4 weeks in the FAS (sows and piglets), 8 weeks in the PWS and 16 weeks in the GS and FIS. Fattening pigs go through FAS, PWS and FIS before to head for slaughterhouse at the age of 28 weeks (main document, Fig 1). Sows' batches can go through GS and FAS several times. Initialization of farms' herd is based on their category, and on information from BDPORC about the herd size. 4-week periodicity of farms is shifted, such as the time frame is not the same for every farm.

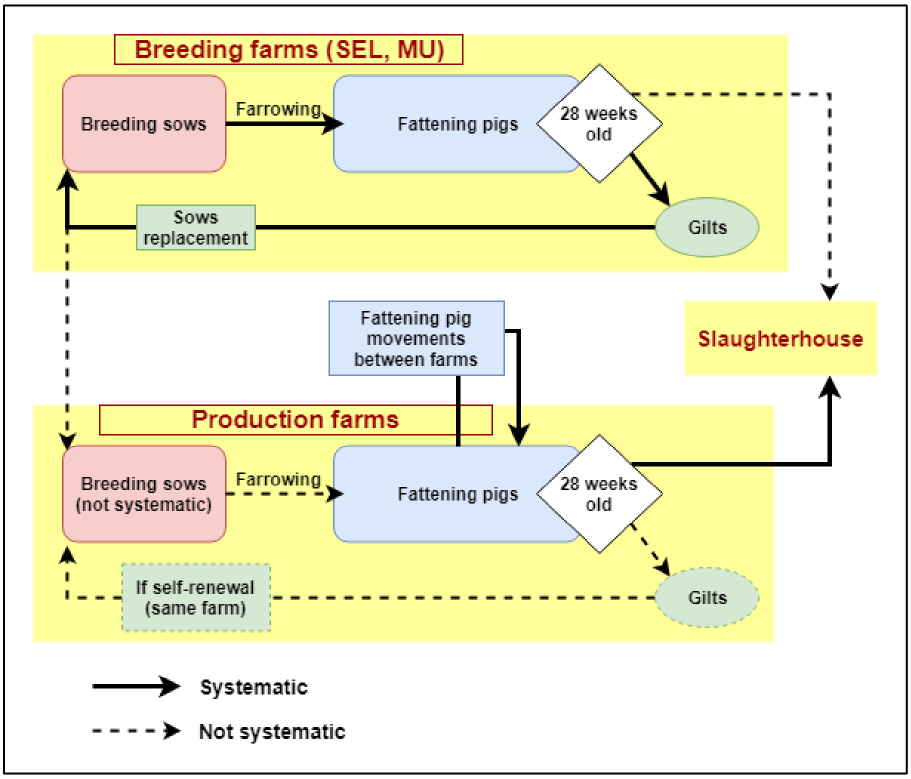


***Figure S1.*** *General overview of the model, with two types of farms (breeding farms and production farms), and two types of herds (breeding and fattening herd).*

3) Pig exchanges between farms

Adjacency matrixes of movements based on BDPORC were filtered in accordance with farms' categories (Table S2). Figure S2 shows the contact matrixes between farm categories. This contact matrix matches data published by the French pig industry on piglets' movements between categories of farms [3].

Every 4 weeks, 3.19% of sows' herd is replaced (based on 41.5% of a sows' herd replaced per year in 2014 [4]), chosen randomly in the batch coming from the FAS and about to enter the GS. They are replaced by gilts from either another farm, according to BDPORC network data, or 28 week-old female pigs of the same farm, when the farm does not appear to import gilts from another farm in the database.

Exportation of fattening pigs occurs either from the FAS room of the exporting farm to the PWS1 room of the receiving farm, or from the PWS2 room of the exporting farm to the FIS1 room of the receiving farm. Exportation times and percentage of pigs from a batch that are exported depend on the exporting farm's category and on BDPORC data (Table S2). Pigs were assumed not to change MRSA status during their transportation from a farm to another.

***Table S2.*** *Rules for production pigs exchanges in the demographic process, depending of the farms' categories.* $P_{exp,farm,n}$ *is the proportion of a batch (compartment) that is exported by a farm at the n-th exportation period (n=1 after Farrowing, n=2 after Post-Weaning). For some categories, we set* $P_{exp,farm,n}$ *to 0 or 1. When not set to 0 or 1,* $P_{exp,farm,n}$ *is calculated as follows:*

$$P_{exp,farm,n} = \frac{Number\ of\ pigs\ of\ a\ batch\ exported}{Mean\ number\ of\ pigs\ in\ a\ batch} = \frac{(Number\ of\ pigs\ exported\ during\ year\ 2014)/13}{(Number\ of\ fattening\ pigs\ in\ the\ farm\ in\ 2014)/(Number\ of\ batches)}$$

| **Farm category** | | **Production pig exchange step (rational based on Fig 2)** | | | |
|---|---|---|---|---|---|
| | | **Farrowing ➔ Post-Weaning** | | **Post-Weaning ➔ Finishing** | |
| **Abbreviation** | **Category** | **Exporting (EP1)** | **Importing (IP1)** | **Exporting (E2)** | **Importing (I2)** |
| FA | Farrowing | 100% of compartment FA(t) ($P_{exp,farm,1}$=1) | Not allowed | 0 ($P_{exp,farm,2}$=0) | Not allowed |
| FPW | Farrowing-Post-Weaning | 0 ($P_{exp,farm,1}$=0) | Allowed | 100% of compartment PW2(t) ($P_{exp,farm,2}$=1) | Not allowed |
| PW | Post-Weaning | 0 ($P_{exp,farm,1}$=0) | Allowed | 100% of compartment PW2(t) ($P_{exp,farm,2}$=1) | Not allowed |
| PWF | Post-Weaning-Finishing | 0 ($P_{exp,farm,1}$=0) | Allowed | $P_{exp,farm,2}$* PW2(t) | Allowed |
| FI | Finishing | 0 ($P_{exp,farm,1}$=0) | Allowed | 0 ($P_{exp,farm,2}$=0) | Allowed |
| FF | Farrowing-to-Finishing | 0 ($P_{exp,farm,1}$=0) | Allowed | $P_{exp,farm,2}$* PW2(t) | Allowed |
| MU | Multiplier | 0 ($P_{exp,farm,1}$=0) | Not allowed | $P_{exp,farm,2}$* PW2(t) (limited to max = 50%) | Not allowed |
| SEL | Nucleus | 0 ($P_{exp,farm,1}$=0) | Not allowed | $P_{exp,farm,2}$* PW2(t) (limited to max = 50%) | Not allowed |

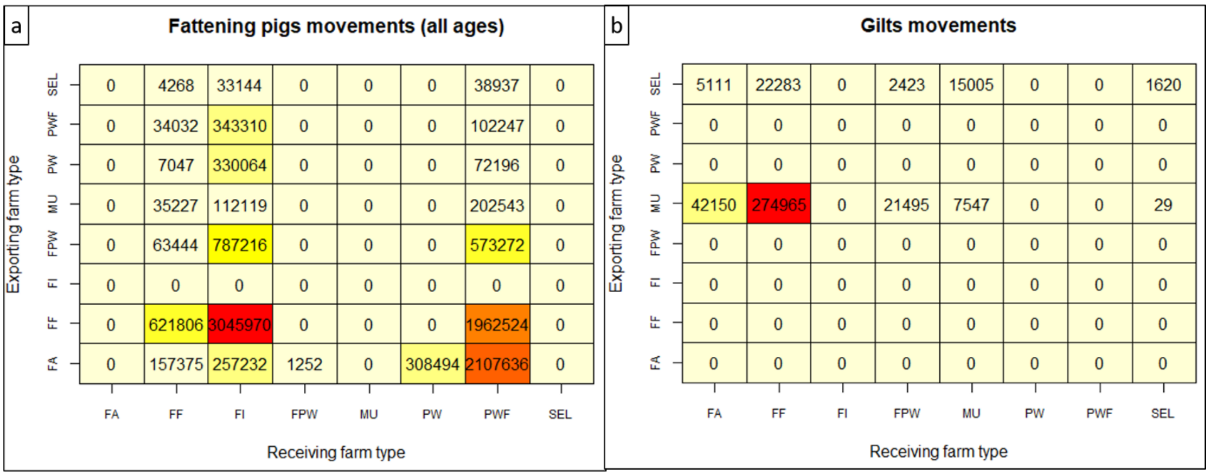


***Figure S2.*** *Contact matrix (number of animals moved in one year) between farm categories, derived from BDPORC data in 2014. a) Fattening pigs movements. b) Gilts movements.*

### S3. Details on the "Invasion paths" method to select sentinel farms

The "Invasion paths" method, was extended from [5] and [6]. When MRSA is introduced in a farm (the seed), it is transmitted from farm to farm. The set of nodes (farms) reached by MRSA constitute the invasion path of the seed farm X, and is written Γ(X). Then, we calculate a Jaccard index $\Theta(X,Y)$ between each pair of seed X and seed Y, as follows:

$$\Theta(X,Y) = \frac{|\,\Gamma(X)\ \cap\ \Gamma(Y)\,|}{|\,\Gamma(X)\ \cup\ \Gamma(Y)\,|}$$

We obtain a network called initial-condition similarity network (ICSN). The nodes are all the possible seeds, and the edges' weights are the value of the Jaccard index Θ between each pair of nodes (between 0 and 1). As in [5], we reduce this network by taking account only the seeds whose invasion path is strictly larger than 4 nodes, to exclude seeds that get artificially a high Jaccard value because their invasion path is only 2 or 3 nodes large, in common with another seed. We can also apply a filter, i.e. removing edges in the ICSN when their weight is lower than a value α ∈ [0,1] [5].

The next step is to calculate the weakly connected components (WCC) of the ICSN. We select randomly one farm in each of the WCC to be a sentinel farm. We repeat iterations for each ICSN to account for this variability (main document, Fig 7). A higher value of filter α makes the ICSN less connected, leading to a higher number of WCC, and then a higher number of sentinel farms.

**S4. Model behaviour under the "realistic" scenario 1**

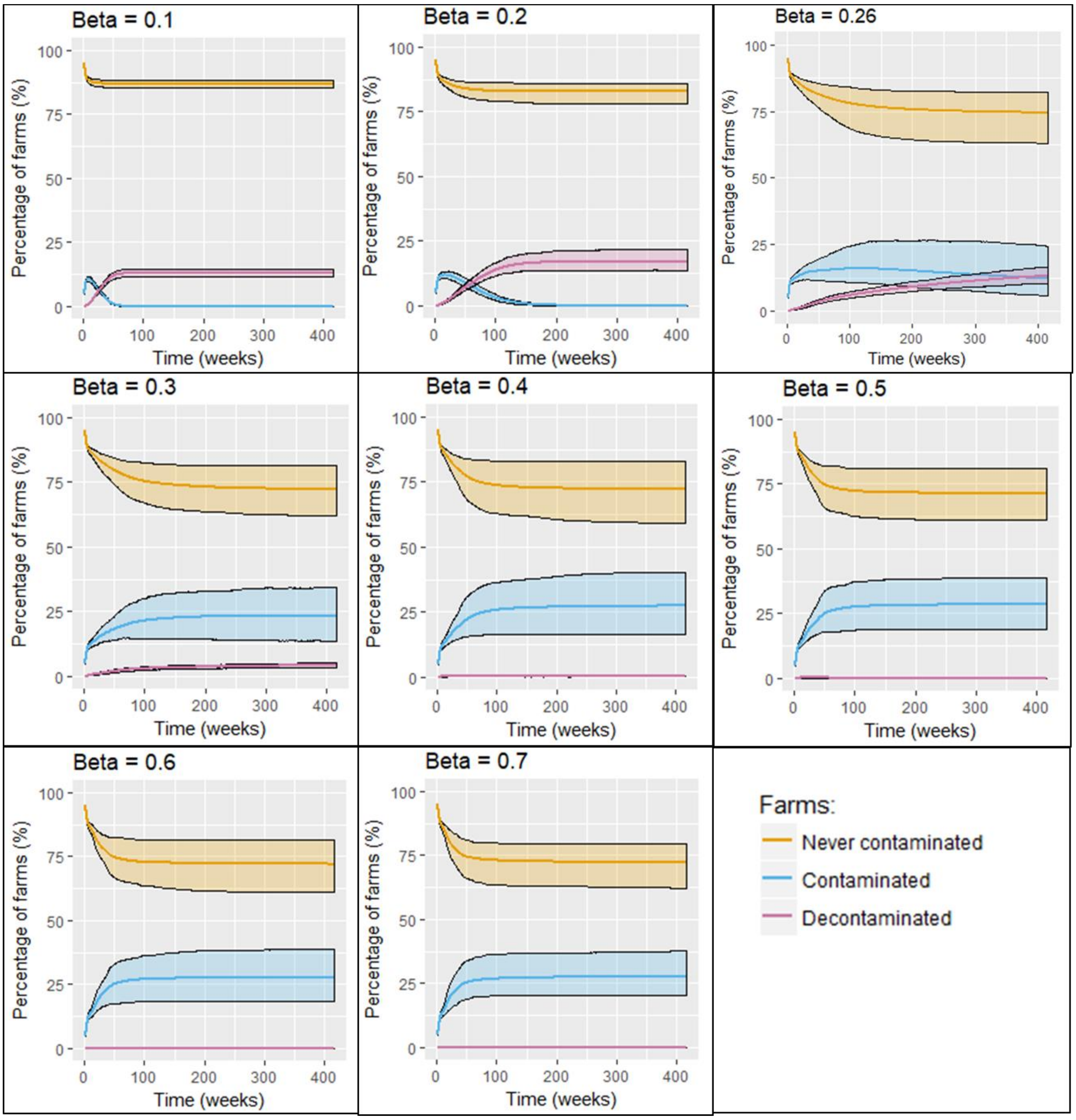


***Figure S3.*** *Under the "realistic" scenario 1, evolution over time of the percentages of farms of the network which have not yet been contaminated by MRSA, of farms which are contaminated, and of farms which have been contaminated but were then decontaminated (local extinctions). Several values of* $\beta$ *were tested. 95% prediction intervals are displayed.*

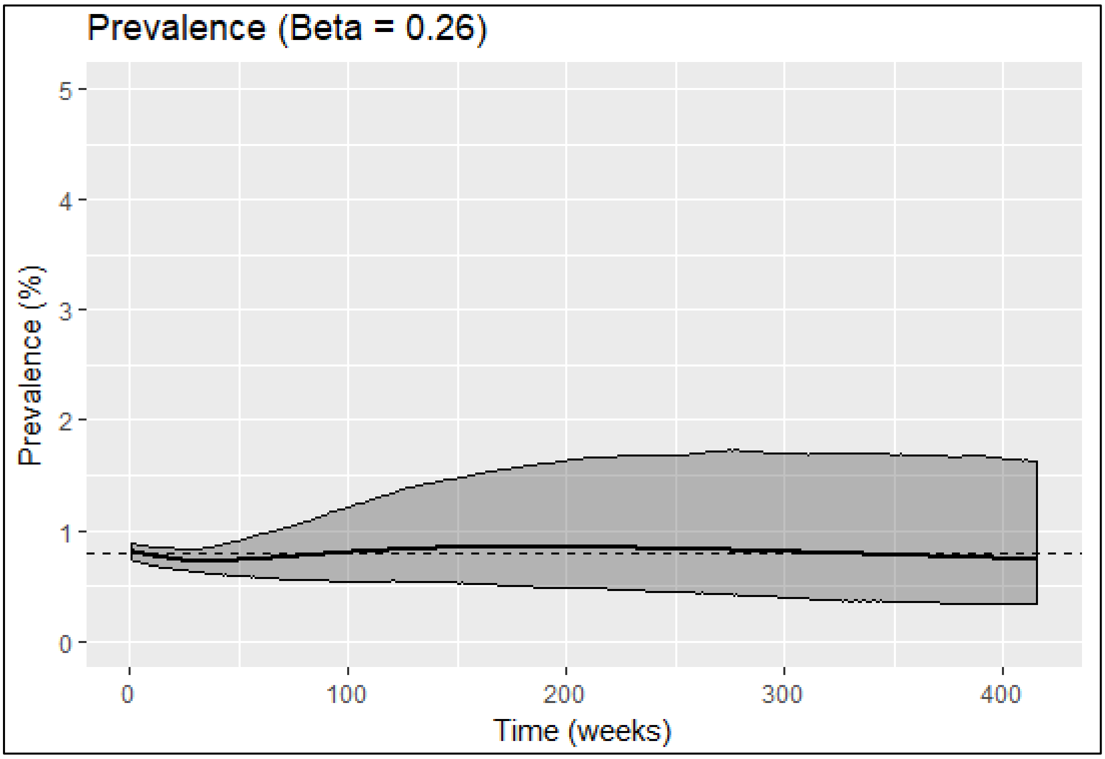


***Figure S4.*** *Under the "realistic" scenario 1, evolution over time (8 years) of the national prevalence in pigs, for β=0.26 (baseline value). Over time, the prevalence is close to the observed carriage of 0.8%* [7,8]*, although it slowly decreases. The 95% prediction interval is displayed.*

**S5. Results of the multivariate analysis to identify factors associated with the spreading potential of seed farms under the "introduction" scenario 2**

***Table S3.*** *Results of the multivariate linear regression. PFC after one year (%) explained by the seed farm's characteristics, in the "introduction" scenario 2.*

| | **Estimate** | **p-value** |
|---|---|---|
| **(Intercept)** | -0.78 | $3.55 \times 10^{-6}$ |
| **Farming category: Breeding farm (Ref: Production farm)** | 2.96 | $< 10^{-15}$ |
| **Size of the production herd** | $1.90 \times 10^{-4}$ | $< 10^{-15}$ |
| **Outdegree** | 0.17 | $< 10^{-15}$ |
| **Outflux** | $5.04 \times 10^{-4}$ | $< 10^{-15}$ |
| **Coreness** | 0.30 | $< 10^{-15}$ |
| **Indegree** | -0.16 | $< 10^{-15}$ |
| **Betweenness** | $6.15 \times 10^{-3}$ | $3.44 \times 10^{-13}$ |
| **Closeness** | $3.94 \times 10^{6}$ | $3.45 \times 10^{-6}$ |

### S6. Additional results – Impact of targeted control measures

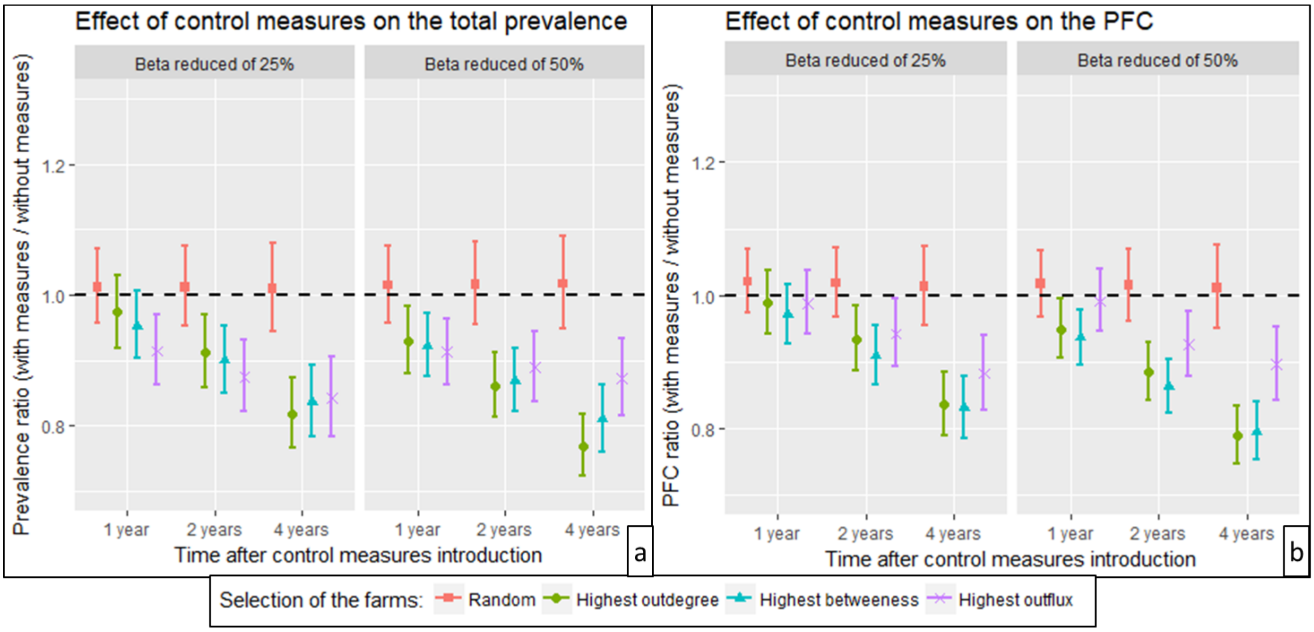


***Figure S5.*** *Impact of control measures implemented in 100 farms selected at random (red dots), in the 100 farms with the highest outdegree (green dots), in the 100 farms with the highest betweenness (blue dots) or in the 100 farms with the highest outflux (purple dots). The ratio "Output value with control measures / Output value without control measures" is depicted at different times (1, 2 and 4 years after control measures implementation) for two model outputs: (a) Total prevalence in pigs and b) Percentage of Farms Contaminated (PFC). "Realistic" scenario 1 was simulated. Two levels of control measure intensity are considered: reducing the transmission parameter β by 25% or 50%. Intervals are Fiellers 95%CI of the t-test for the ratio of two means: the mean values of outputs of 300 model iterations for the case control measures are applied VS the case control measures are not applied.*

### S7. Additional results – Sentinel selection for targeted surveillance

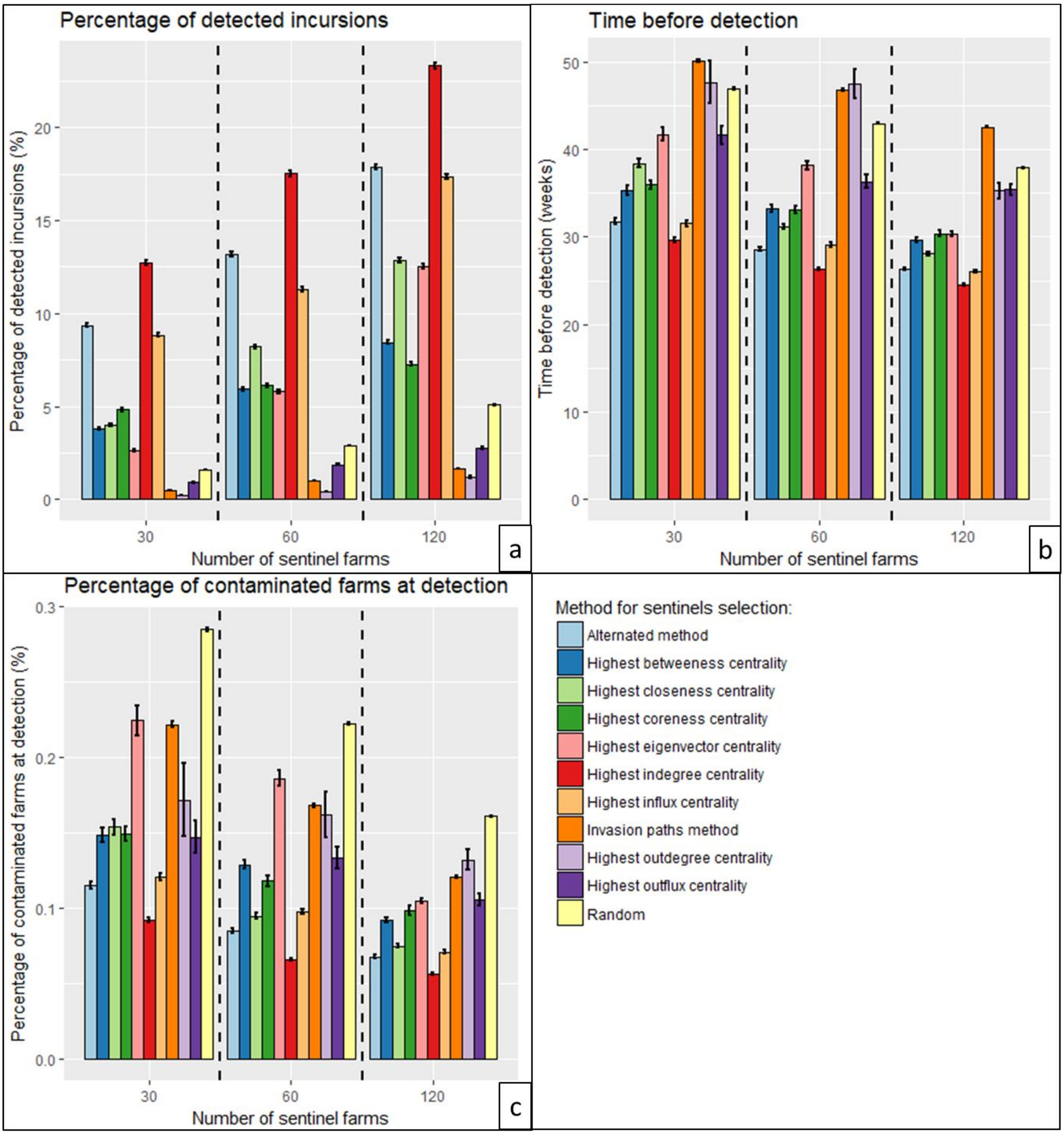


***Figure S6.*** *Comparison of the efficiency of several methods for sentinel selection, based on 3 efficiency criteria: Percentage of MRSA outbreaks detected (a), Time before detection (b), and Percentage of farms contaminated (PFC) at detection (c). Several sizes of sentinel sets were tested: 30, 60 and 120 sentinel farms. For each set of sentinel farms, the surveillance performance was assessed based on the average of model simulations of MRSA introduction starting from all farms housing breeding sows in the network. Intervals are 95%CI of the t-test. 100 different sets of sentinels were considered with the "Random" and "Invasion paths" methods (see definitions of network centrality indicators in SM1).*
